# Destabilization of the holo-DNA Polδ by loss of Pol32 confers conditional lethality that can be suppressed by stabilizing Pol31-Pol3 interaction

**DOI:** 10.1101/2021.02.11.430699

**Authors:** Kenji Shimada, Monika Tsai-Pflugfelder, Niloofar Davoodi Vijeh Motlagh, Neda Delgoshaie, Jeannette Fuchs, Susan M. Gasser

## Abstract

DNA Polymerase δ plays an essential role in genome replication and in the preservation of genome integrity. In *S. cerevisiae*, Polδ consists of three subunits: Pol3 (the catalytic subunit), Pol31 and Pol32. We have constructed *pol31* mutants by alanine substitution at conserved amino acids, and identified three alleles that do not confer any disadvantage on their own, but which suppress the cold-, HU- and MMS-hypersensitivity of yeast strains lacking Pol32. We have shown that Pol31 and Pol32 are both involved in translesion synthesis, error-free bypass synthesis, and in preservation of replication fork stability under conditions of HU arrest. We identified a solvent exposed loop in Pol31 defined by two alanine substitutions at T415 and W417. Whereas pol31-T415A compromises polymerase stability at stalled forks, *pol31*-W417A is able to suppress many, but not all, of the phenotypes arising from *pol32*Δ. ChIP analyses showed that the absence of Pol32 destabilizes Polε and Polα at stalled replication forks, but does not interfere with checkpoint kinase activation. We show that the Pol31-W417A-mediated suppression of replication-stress sensitivity in *pol32*Δ stems from enhanced interaction between Pol3 and Pol31, which stabilizes a functional Polδ.

## Introduction

DNA Polymerase δ (Polδ) cooperates with DNA Polymerase *α* to synthesize the lagging strand during genomic replication, yet it also plays essential roles in leading strand synthesis and in many DNA repair pathways(Zhou, Lujan et al., 2019) (reviewed in(Prindle & Loeb, 2012)). In the model organism *S. cerevisiae*, Polδ consists of three subunits: the catalytic subunit Pol3, and the regulatory subunits, Pol31 and Pol32(Gerik, Li et al., 1998). The genes encoding Pol3 and Pol31 are essential, while *POL32* is non-essential. Nonetheless, pol32-deficient strains are compromised for break-induced replication, a recombination event that allows restart at collapsed replication forks(Hanna, Ball et al., 2007, Lydeard, Jain et al., 2007) and show sensitivity to alkylating agents (e.g., methyl methanesulfonate or MMS) and to dNTP depletion by hydroxyurea (HU), which both perturb replication fork progression(Vijeh Motlagh, Seki et al., 2006). Like the budding yeast enzyme, mammalian DNA Polδ is constituted of subunits POLD1-D3, but it harbours a smaller subunit, POLD4, that appears to serve as an inhibitor of DNA Polδ activity (Zhang, Zhao et al., 2013, Zhang, Zhou et al., 2007). This fourth subunit was initially detected in fission yeast (*Sp* Cdm1), and unlike the other subunits of Polδ, this subunit is not essential for growth (Zuo, Gibbs et al., 1997).

The roles played by the second and third subunits of Polδ, Pol31 and Pol32, and their distinct contributions to holoenzyme function are still unclear. A purely structural function, that of linking the catalytic Pol3 to the third subunit has been proposed for Pol31. That function alone would not explain, however, why *POL31* is an essential gene, while *POL32* is not. Here we characterize the functions of the Polδ second subunit, Pol31, at replication forks under DNA damage stress, and its role in compensating for the loss of the nonessential subunit Pol32. Both Pol31 and Pol32 are of interest for cancer biology, as the human counterparts POLD2 and POLD3 are commonly found amplified in tumours (Fuchs et al., in press; {Beroukhim, 2010 #2457}).review; (Beroukhim, Mermel et al., 2010)).

Human cancer cells are characterized by a persistent replication stress, which arises from inappropriate origin firing and a failure to coordinate replication with transcription (refs). Consistent with genetic studies in yeast and flies which indicate important roles in the maintenance of genome stability during replicative stress, human Polδ subunits are frequently overexpressed in a wide range of human cancers, while POLD4 is occasionally downregulated or lost (Fuchs et al., in press). Whereas POLD2 and POLD3 are often highly expressed, the catalytic subunit POLD1 acquires point mutations that result in a defective proofreading activity for the polymerase. POLD1 mutations have been associated with increased tumor incidence and decreased survival in mouse models {Goldsby, 2001 #2472}, and both germline and sporadic mutations in the *POLD1* proofreading exonuclease domain were found in human cancers (Rayner et al.,.2016, Palles et al., 2013; Nicolas et al., 2016, Zhang et al., 2020) and were linked with a hypermutator phenotype.

Human cancer cells are characterized by a persistent replication stress, which arises from inappropriate origin firing and a failure to coordinate replication with transcription(Macheret & Halazonetis, 2018). Consistent with genetic studies in yeast and flies which indicate important roles in the maintenance of genome stability during replicative stress, all four Polδ subunits (POLD1 - POLD4) are frequently overexpressed in human cancers. In addition, the catalytic subunit POLD1 acquires point mutations that result in a defective proofreading activity for the polymerase, which has been associated with increased tumor incidence and decreased survival in mouse models (Goldsby, Lawrence et al., 2001). Indeed, both germline and sporadic mutations in the *POLD1* proofreading exonuclease domain have been identified in a range of human cancers (Rayner, et al., 2016).

A study in U20S cells demonstrated that the POLD3 and POLD4 subunits of Polδ are required for DNA synthesis and cell cycle progression in the presence of replication stress induced by cyclin E overexpression. This study also demonstrated that POLD3/POLD4 accounted for a significant portion of genomic copy-number alternations (CNA) generated under prolonged conditions of oncogene-induced replication stress. Based on these findings the authors proposed that BIR repair of compromised replication forks by POLD3 may partly account for genomic duplications in human cancers (Costantino, et al., 2014). However, it is not clear whether Polδ’s role of in replication stress is restricted to BIR-mediated restart. Indeed, Polδ has also been shown to be involved in translesion synthesis (Hirota, et al., 2016).

Whereas the deletion of Pol32 in budding yeast is not lethal in the absence of damage, it is synthetic lethal with many factors involved in replication checkpoint response, namely, the reduction of Mec1(ATR) checkpoint kinase activity, Mrc1, the RecQ helicase Sgs1 and Mre11 (Tong, et al., 2001) reviewed In Hustedt et al., 2013). Moreover, it shows a strong conditional lethality on HU with the loss of 9-1-1 complex (Rad17, Ddc1, Mec3) and its loader, Rad24, both of which contribute to the activation of the Mec1-Rad53 checkpoint cascade on HU (Hustedt et al., 2015). Finally, *pol32Δ* is also lethal with a number of chromatin factors required for DSB repair in S phase, as well as the E2 Ubiquitin ligase Ubc13, on 100mM HU (Hanna et al., 2007, Karras & Jentsch, 2010). Because the deletion of the gene encoding the second subunit of Pol δ, Pol31 (POLD2) is lethal in yeast, it has been characterized less thoroughly.

In an attempt to understand the function of Pol31 in detail, an alanine-scanning mutagenesis of the most highly conserved amino acids within the protein’s ten subdomains (Reynolds & MacNeill, 1999) was carried out (Vijeh Motlagh et al., 2006). Although most alanine substitutions did not confer overt phenotypes, six novel temperature-sensitive (ts) or cold-sensitive (cs) *pol31* alleles were isolated, which mapped to the conserved regions III, IV, VII, VIII or IX in the linear structure (Vijeh Motlagh et al., 2006). The deletion of genes encoding fork stabilizing factors, namely *SGS1, RAD52, SRS2, MRC1* or *RAD24*, was deleterious in combination with the temperature-sensitive *pol31* alleles, but not with those that were temperature-permissive (Vijeh Motlagh et al., 2006). This argues for a requirement for recombination and checkpoint functions in processing the DNA lesions or structures that form as a consequence of replication by a defective Polδ. Intriguingly, all conditional *pol31* alleles, regardless of their phenotype as a single mutant, became dependent on the presence of an intact Pol32 subunit, suggesting that the two smaller subunits of the Polδ complex complement partially within the complex.

In addition to these ts- and cs- alleles, three specific temperature-permissive *pol31* alleles (pol31-D297A, -W417A, and -EF463-464AA, hereafter *pol31-D297*, W417, and -EF463) were identified as suppressors of the cold-, HU- and MMS-sensitivities of *pol32Δ* cells in the alanine-scanning mutagenesis. We have pursued their mode of action, because by understanding the gain-of-function arising from these mutations, we should be able to identify the functions provided by Pol32, that Pol31 normally cannot supply. None of the pol32*Δ*-suppressor mutations mapped to the interface of Pol31 with Pol32, nor to the interface with PCNA. Rather we find that the *pol31* allele W417 enhances the interaction with Pol3, while an adjacent cold-sensitive allele, T415, compromises this binding. These two amino acids thus identify a key interface of Pol31 with the C terminal Fe-S cluster of Pol3. The same mutations also affect interaction with the conserved C-terminal Zn-finger in the translesion polymerase (TLS) Rev3 (polζ; (Makarova, Stodola et al., 2012)). We show that enhanced Pol31-Pol3 interaction correlates with an improved ability to maintain replication fork stability and resume replication after prolonged challenge by HU. Moreover, the DNA Polδ holoenzyme is stabilized by enhanced interaction between Pol31 and Pol3, even in the absence of Pol32. Finally, after prolonged exposure to HU in *pol32Δ* cells, *we* find that DNA polymerase α and ε also become unstable at stalled forks, and that this too is suppressed in part by enhancing the affinity of Pol31 for Pol3. This argues for crosstalk among replicative polymerases at the stalled fork. Overall levels of Pol3 are elevated in the W417 mutant and reduced by T415, showing that holoenzyme complex abundance depends on stable inter-subunit interactions. In conclusion, we propose that if one of the other smaller subunits of the holo-Polδ is compromised, the level of catalytic enzyme drops, and improved interaction of the other two remaining subunits is able to restore DNA Polδ function. This provides an explanation of the suppression conferred by the *pol31*-W417 mutation as well as for the persistent upregulation of these subunits in cancer cells, which experience persistent replication stress.

## Results

Based on protein sequence comparisons of the POLD2/Pol31 subunit across eukaryotes, ten conserved regions (I-X) were identified and the residues with the highest conservation were mutated one by one to alanine (Reynolds & MacNeill, 1999). Three alanine substitutions in Pol31, namely, D297, W417 and EF463, which respectively map to the domains VI, VIII and X, showed no sensitivity to either HU, MMS, nor to growth at low or high temperature (Figure 1A,B). This is in contrast to *pol32A* Based on the crystal structure of the human orthologue of Pol31-Pol32_N-term_ (Baranovskiy, Babayeva et al., 2008), we built a homology model of Pol31, and indicated the positions of the novel *pol31* mutations on the model. The three alleles do not cluster in one domain or interface, although the D297 and EF463 substitutions are not far from the Pol32 binding surface (Figure 1C). In contrast, the third suppressor mutation (pol31-W417) mapped to a solvent exposed loop on the surface of Pol31, in a surface patch that has the highest conservation score in Pol31. Intriguingly, an alanine substitution at T415 conferred HU, MMS and cold sensitivity in an otherwise wild-type strain (Figure 1B). Thus, the T415 and W417 mutations identify a functional domain at which the T to A substitution at aa 415 compromises function, and the W to A substitution at aa 417 suppressed insufficiencies provoked by *pol32Δ*.

**Figure 1:**
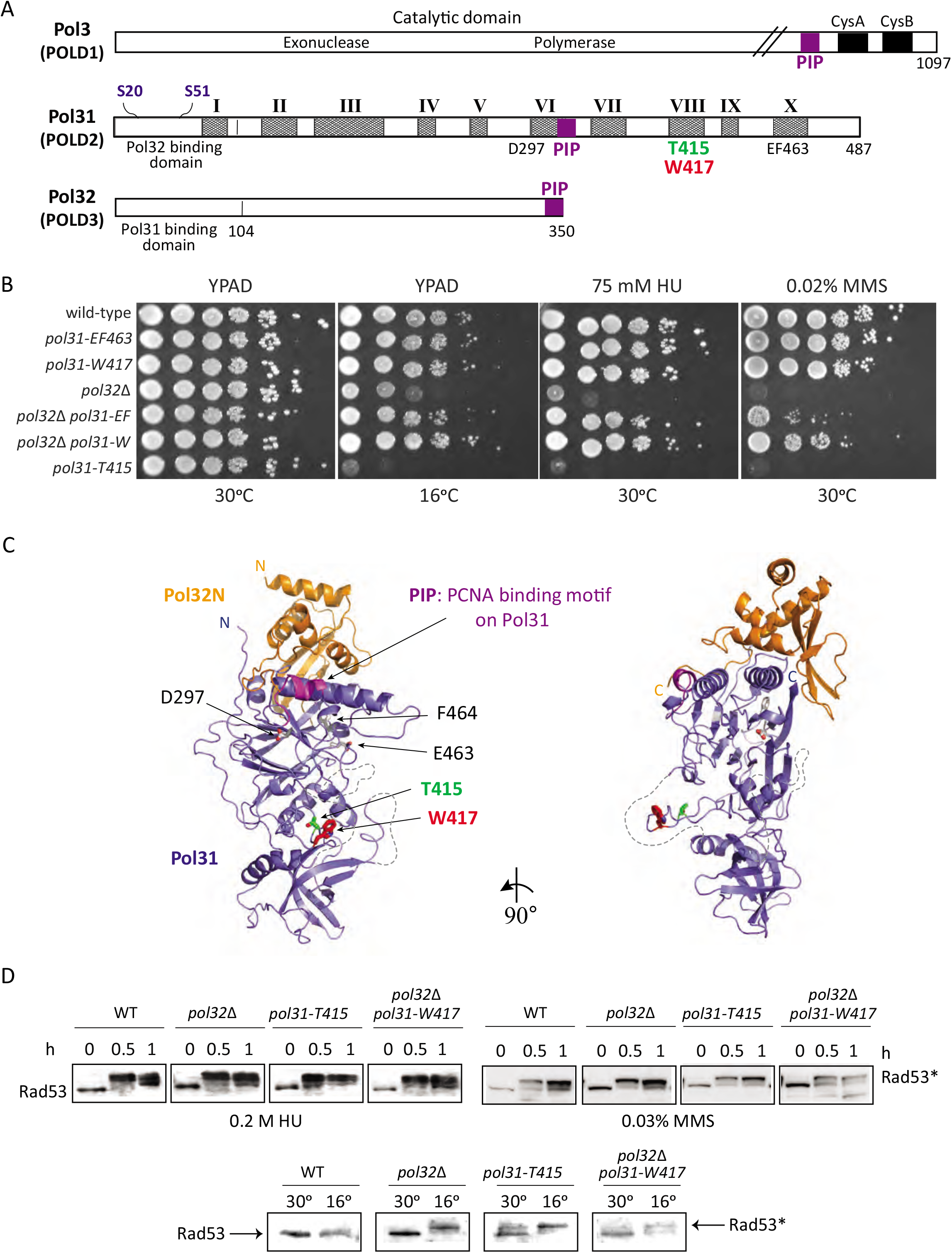
Mapping of the *pol31* mutants that suppress the HU-, MMS- and cold sensitivity of *pol32Δ* as well as *pol31-T415* mutant which exhibit HU-, MMS- and cold sensitivity. A) Linear map of the three subunits of DNA Polδ: Pol3, Pol31 and Pol32, with interacting domains, conserved domain of Pol31 (conserved regions, shaded) and indication of the mutants used in this study. PIP: PCNA interaction sites. B) Serial dilution (10x series) drop assays demonstrate the sensitivities of the pol31-T415 and *pol32Δ* mutants, and the ability of *W417-pol31*, and EF463-pol31 mutants to suppress the lethality of *pol32Δ on 75 mM HU or 0.02%* MMS or at 16°C. C) Homology model of the Pol31-Pol32N complex. Pol31 (blue) and Pol32N (orange) are displayed as cartoon models in two orientations rotated around a vertical axis by 90°. Specific residues which were mutated in this study are shown as sticks in atom colors (T415, green; W417, red; D297, E463-F464, grey). The Pol31 PCNA binding motif is highlighted (magenta) and N- and C-termini are labeled. Flexible loops which were not included in the homology modeling calculation (see Materials and Methods) are shown as dashed grey lines. (D) Rad53 phosphorylation upshift (Rad53*) monitors checkpoint kinase activation after treatment with HU or MMS at 30°C, or incubation at 16°C, which causes lethality in *pol32Δ* as well as *pol31*-T415 mutants. Checkpoint activation is intact in *pol32Δ, pol31-T415*, and *pol32Δ, pol31-W417* double-mutants after treatment with HU, MMS or during incubation at 16°C as seen by Rad53 upshift. Rad53 upshift in *pol32Δ* mutant at low temperature is partially suppressed by additional mutation of *pol31-W417*.

*We* can rule out that the failed growth arises from impaired checkpoint activation, because Rad53 (CHK2) phosphorylation after HU treatment is intact in *pol32Δ* as well as in *pol31-T415*, and the activation still occurs in the pol31-W417 *pol32Δ* double mutant (Figure 1D). Intriguingly, we note that growth at 16°C alone can trigger checkpoint activation in both *pol32Δ* and the pol31-T415 mutants, although the exact mechanism is unclear. The cold-sensitive activation of Rad53 in cells lacking Pol32 is partially suppressed by the pol31-W417, consistent with the improved growth of the pol31-W417 *pol32Δ* double mutant on 75 mM HU (Figure 1B,D).

### The pol31-W417 mutation improves Pol3-Pol31 binding independently of Pol32, *while* pol31-*T415 compromises it*

We examined the involvement of the exposed loop in Pol31 in protein-protein interactions, by yeast two-hybrid and pull-down assays with its most likely interaction partner, the Pol3 C-terminal domain (CTD). The C-terminal domain of Pol3 contains two metal-chelating Zn-finger like motifs called CysA and CysB, the former being a Zn-finger involved in PCNA (Garcia et al., 2004). The C-terminal domain of Pol3 contains two metal-chelating Zn-finger like motifs called CysA and CysB, the former being a Zn-finger involved in PCNA (Acharya et al., 2011, Khandagale et al., 2020) binding, while the latter is a Fe-S center (Jain et al., 2019). Interestingly, quantitative two-hybrid assays using CysA and CysB domains of Pol3 with full length Pol31 show that the pol31-T415 mutation compromises the interaction of Pol31 with the Pol3-CTD while the *pol31-W417* mutation does not (Figure 2A). Moreover, when we use only the CysB domain of Pol3, which is suggested to be important in Pol31 binding (Jain et al., 2019), we find that while pol31-T415 mutation compromises the interaction of Pol31 with the Pol3-CysB, the *pol31-W417* mutation actually increases the affinity to Pol3-CysB, while *pol31-T415* mutation abolishes the interaction (Figure 2B). Importantly, this was true both in the presence and in the absence of Pol32, confirming that the effect of the mutations are direct, not reflecting an improved Pol31-Pol32 interaction that might stabilise Pol3 binding indirectly.

**Figure 2:**
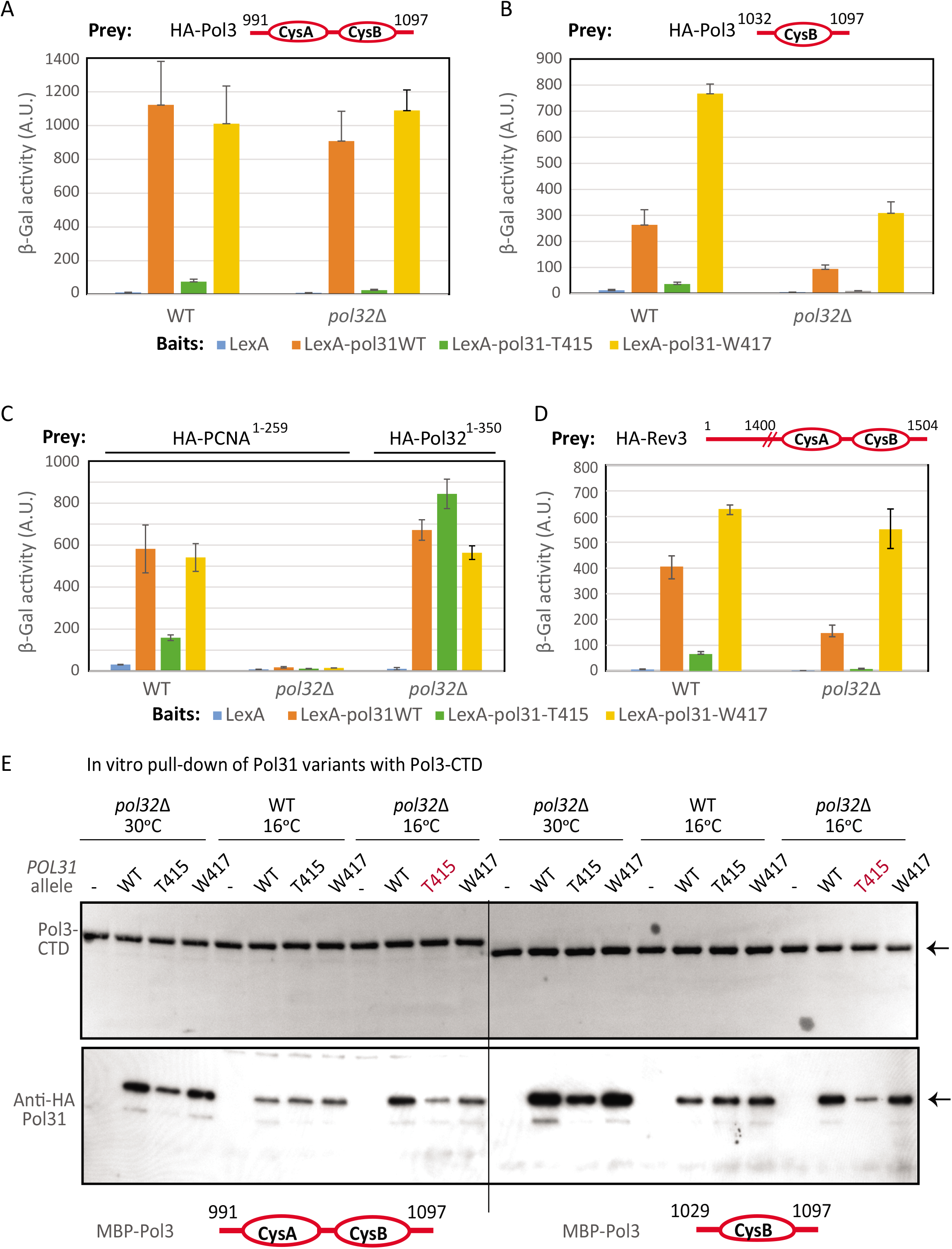
Pol31 interacts with the C-terminal domain of DNA Pol3, and is sensitive to W417 and T415 mutations. A) Yeast two-hybrid interaction performed by standard methods is quantified by βgalactosidease units in solution (A.U.) which are expressed linearly based on the strength of bait and prey interaction. The assay is performed in the indicated backgrounds (wild-type or *pol32Δ*) with the indicated bait and prey constructs. We find that irrespective of the presence of Pol32, Pol31 wildtype and *pol31-W417* mutant bind to the Pol3 C-terminuS_(991.1097)_ bearing the paired CysA and CysB domains, but this interaction is compromised by the *pol31-T415* mutation. B) As in A), but the yeast two hybrid assay is performed using a prey that contains only the more distal CysB FE-S cluster of the Pol3 C-terminus _(1032-1097)_. The interaction is again compromised by the *P¤I3I-Ĩ4I5* mutation. Although the interaction between Pol3-Cys B domain and Pol31 is slightly weakened in the absence of Pol32, *pol31-W417* mutation actually increases the affinity of this binding, irrespective of the presence of Pol32. C) Yeast two-hybrid assays showing Pol31 wild-type and Pol31 mutants interaction with PCNA in a manner dependent on Pol32. *pol31-T4l5* reduces interactions with PCNA, but *pol31-W417* does not. All Pol31 interactions with PCNA are lost in the absence of Pol32. Similar assays show Pol31 binds the Pol32 N-terminus, independently of endogenous Pol32 and *pol31* mutations. D) As in C), Yeast two-hybrid assays performed as above with the indicated baits and prey. This assay shows that the *pol31-T415* mutation strongly reduces binding affinity to Rev3 (the catalytic subunit of Polζ), while *pol31-W417* enhances Rev3 binding. Tis is independent of endogenous Pol32. Still the loss of Pol32 partially weakens interaction between wild-type Pol31 and Rev3. E) Pull down assays for Pol31 and variants expressed in yeast extracts by recombinant, purified MBP-Pol3 C-terminal domains. The scheme for the pulldown assay is in Supplemental figure 1. The cs mutant *pol31-T4l5* shows reduced binding interaction with Pol3-_991.1097_ and with Pol3-_1032-1097_ as compared to the wild-type *POL31*. The reduced binding is detected only in cell extracts depleted for Pol32 (*pol32Δ* irrespective of growth temperature. -: vector only, WT: expression of full length HA-tagged Pol31, T415: expression of full length HA-tagged Pol31-T4l5 mutant; W417: expression of full length HA-tagged Pol31-W417 (HA-tagged Pol31 is 78kDa)

Pol31 has also been implicated in tethering the holo-Polδ to PCNA (Acharya et al., 2011), thus we checked the impact of the Pol31 mutations on PCNA interaction. Again there is a drop in Pol31-PCNA binding when T415 is mutated, which is not present in W417. However, the interaction of Pol31 with PCNA by two-hybrid is completely dependent on the presence of endogenous Pol32 (Figure 2C). An altered binding to PCNA thus cannot explain suppression of *pol32Δ* phenotypes, since the Pol31-PCNA interaction requires Pol32. Finally, we showed that neither mutation compromises the interaction of Pol31 with the Pol32 N-terminus, a result predicted by the published crystal structure ((Johansson, Garg et al., 2004); Figure 2C). We conclude that the cs phenotypes of *pol31*-T415 and the suppressor function of *pol31-W417*, may be acting through a reduction and enhancement, respectively, of Pol31 binding to the catalytic subunit Pol3. Intriguingly, however, the pol31-T415 and *pol31-W417* mutations have the same differential effects on the interaction of Pol31 and the CTD of the translesion synthesis polymerase, Rev3 (Pol ζ). This interaction and the impact of the Pol31 alleles on Rev3 binding are independent of Pol32, as in the case of Pol3 (Figure 2D).

We confirmed the two hybrid results with an *in vitro* pull-down assay, using a HA-tagged Pol31 in whole yeast cell extracts, recovered on beads bound to recombinant MBP-Pol3-CTD, which was expressed and purified from *E. coli* (see Materials and Methods). We see the same reduced interaction of the Pol3-CTD with HA-pol31-T415, while there is little or no impact of the HA-pol31-W417 mutation on Pol3-CTD binding, at least in *pol32Δ* extracts (Figure 2E). We find that these fluctuations are masked in the presence of Pol32, which suggests that in wild-type extracts a trimeric complex (Pol3-Pol31-Pol32) is formed, and that the Pol32-Pol3 interaction masks the effect of the pol31-T415 mutation on protein-protein interactions, at least as monitored in cell extracts.

### pol31-W417 can suppress the MMS sensitivity of pol32Δ through PRR pathways without the TLS polymerase, Rev3

The binding of Pol31 to both Pol3 and Rev3, and the modulation of this interaction by the T415 and W417 mutations may account for the sensitivity of the former allele to MMS and HU, and the ability of *pol31-W417* to suppress the sensitivity of *pol32Δ* to growth at 16°C, HU and MMS. In recovery assays, in which the cells synchronized in G1 are released into MMS, the cs mutant *pol31*-T415 strongly impaired cell survival, almost as profoundly as *pol32Δ*, while *pol31-W417* was able to suppress the MMS sensitivity of a strain lacking Pol32, and recover on MMS similar to wild-type yeast (Fig. 3A). There are two pathways used by yeast cells undergoing replication to repair alkylated bases, and these both can occur after the passage of the replicative polymerases (so-called post-replicative repair (PRR) pathways). The first is translesion synthesis (TLS) which involves the recruitment of error-prone translesion polymerases like Polζ (Rev3), while the second is largely error-free, and involves bypassing the mutation by copying the intact strand, through an exchange that requires Rad5, Mms2 and Ubc13 (Figure 3B). Key to pathway choice is the ubiquitination status of PCNA. TLS requires the covalent mono-ubiquitination of K164 of PCNA which is catalyzed by a complex comprised of Rad6 and Rad18 proteins, the E2 and Ring-finger E3 ligases, respectively (Carlile, Pickart et al., 2009, Gangavarapu, Haracska et al., 2006, Halas, Krawczyk et al., 2016, Xu, Blackwell et al., 2015). The poly-ubiquitination of the same residue on PCNA (K164), on the other hand, shunts the lesion to the error-free bypass pathway, triggered by Mms2, Rad5 and Ubc13. PCNA poly-ubiquitination activates a recombination-like DNA damage-avoidance mechanism.

**Figure 3.**
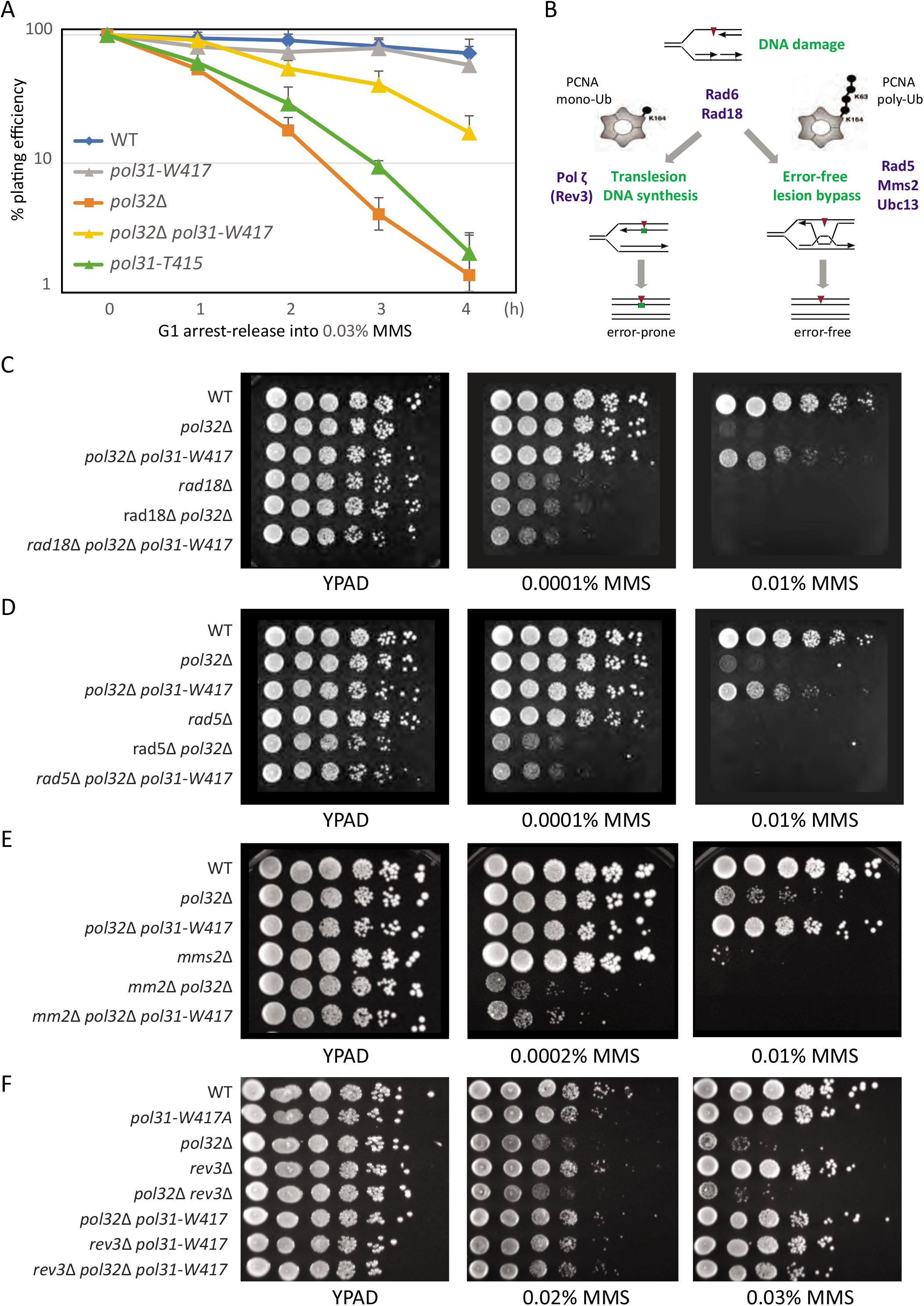
Survival on MMS requires Pol32, and pol31-W417 suppression acts through the Error-free lesion bypass pathway. A) Quantitative recovery assay showing that the *pol31-W417* mutation suppresses the growth defects of *pol32Δ* in liquid culture on MMS. Cells were G1-arrested by α-factor, released into 0.03% MMS, sampled at indicated time-points and plated on YPAD plates. Colonies are scored after 3 days. B) Post replication repair (PRR) pathways and key regulatory proteins. C) Serial dilution (10x) drop-assay showing *rad18Δ* combined with *pol31-W417* and *pol32Δ* mutants on YPAD plates with or without the indicated concentration of fresh MMS. D) as C, but for the *rad5Δ* mutant, The *rad18Δ* and *rad5Δ* mutants are very sensitive to 0.01% MMS on their own and dead at 0.02% MMS. They are synthetic lethal in combination with *pol32Δ*, on 0.01% MMS, *pol31*-W417 can’t suppress this. E) as C, but with of *mms2Δ* showing synthetic lethality in combination with *pol32Δ* on 0.01% MMS, *pol31-W417* can’t suppress this. F) as C, with *rev3Δ* mutant. The *rev3Δ* mutant on its own is not sensitive to 0.02% or 0.03% MMS, but in combination with *pol32Δ* is. *pol31-W417* can still suppress the growth defects of *pol32Δ* i n the absence of rev3*Δ* (Triple mutant).

To see if *pol31-W417* suppression of the *pol32Δ* sensitivity to MMS depends on one or the other PRR branch, we made double mutants with the key regulators of TLS and Error-error free lesion bypass. If W417 suppression acts by enhancing one or the other pathway, we expect to see a loss of suppression in a triple mutant that eliminates the pathway. We started with the upstream regulator, Rad18. Indeed, we see that the ability of *pol31-W417* to suppress the sensitivity of *pol32Δ* to 0.01% MMS is lost in the absence of Rad18 (Figure 3C). The suppression is also lost in the absence of Rad5 and Mms2 (Figure 3D,E), placing pol31-W417’s compensation for the *pol32Δ* MMS sensitivity on the pathway of lesion bypass through strand exchange (Fig. 3B-E). In contrast, the loss of Rev3 did not compromise the suppression of MMS hypersensitivity in the *pol31-W417 pol32Δ* double mutant (Figure 3F). This argues genetically, that Pol31-W417 can act without recruiting Rev3 to suppress the sensitivity of *pol32Δ* to MMS, even though interaction with Rev3 is enhanced by the mutation. The results of the drop assays are confirmed in a more quantitative colony recovery assay, in which cells after G1-synchronization are released into 0.03% MMS for increasing amounts of time and then plated to monitor survival (similar to the experiment in Fig. 3A). Based on acute exposure and recovery, we also see that *pol31-W417* can suppress the sensitivity of *pol32Δ* in the absence of Rev3 (Supplemental Figure S2). We propose that the suppression pathway makes use of the enhanced affinity of Pol31-W417 to Pol3 either through Error-free lesion bypass, or through TLS mediated by Polδ. Indeed, in DT40 cells, DNA Polδ has been implicated as an alternative polymerase for translesion synthesis in DT40 cells (Hirota et al., 2016).

### pol31-W417 can suppress the HU sensitivity of *pol32Δ* independently of PRR regulation

Yeast cells deficient for Pol32, or bearing the pol31-T415 mutation, are not only sensitive to MMS, but also to HU (see Figure 1B). In addition to the drop assay shown in Figure 1B, we carried out an assay that monitors the ability of cells to recover from acute fork arrest by dNTP depletion, that is, to form colonies after a transient exposure to 0.2M HU. We find that pol31-T415 strains are even more sensitive to HU than a *pol32Δ* strain, and that *pol31-W417* is able to partially suppress this defect in a *pol32Δ* strain (Figure 4A). The suppression is even more pronounced when cells are exposed to both MMS and HU (Figure 4B).

**Figure 4.**
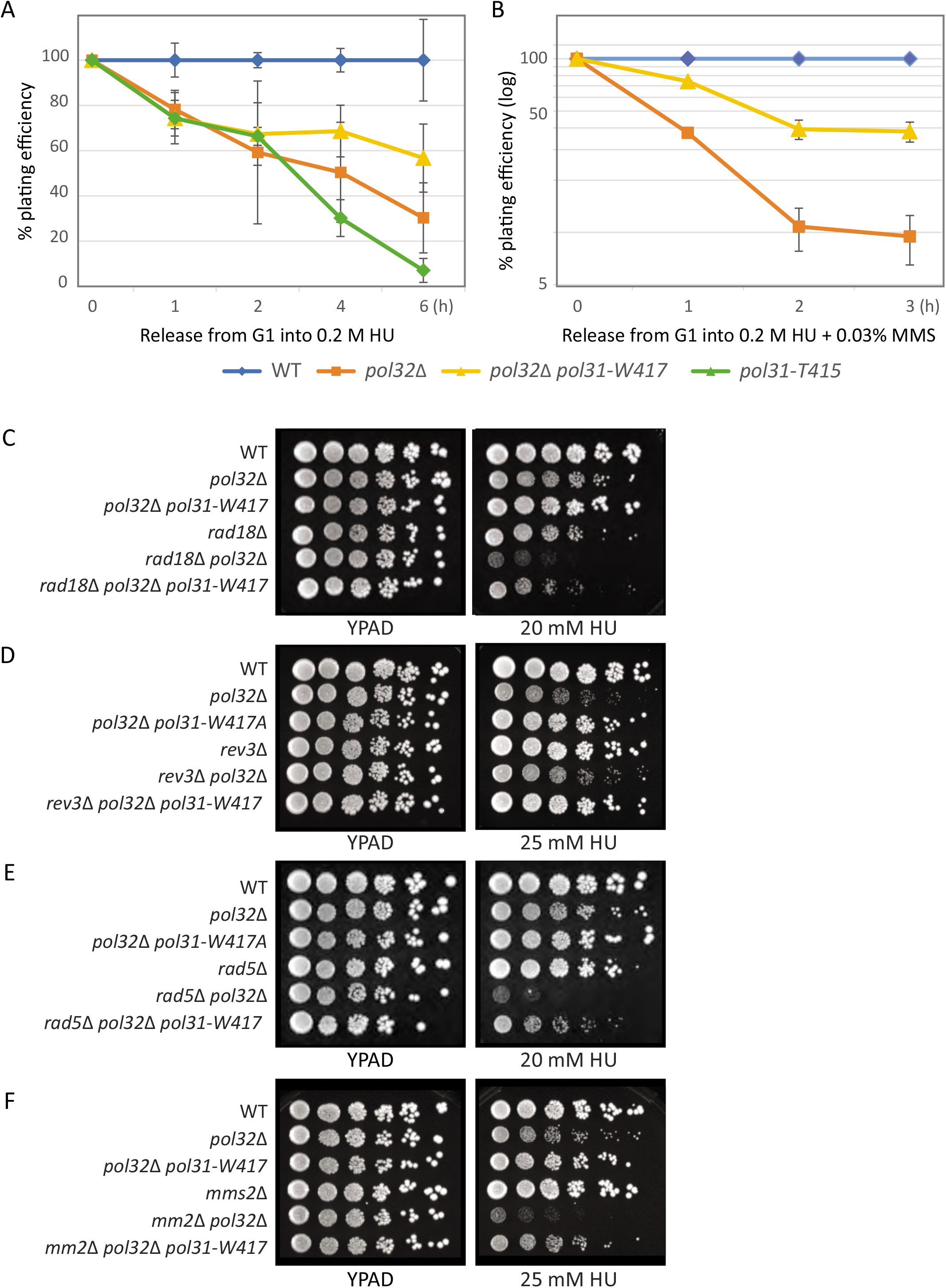
Survival pathway analysis on HU: pol31-W417 suppresses defects of pol32Δ. A) Colony outgrowth assay for exposure to acute levels of HU for the indicated times. Isogenic strains bearing the indicated mutations were synchronized in G1 and released into 0.2M HU for 1 to 6 h prior to plating on YPAD. Colony scoring was carried out in triplicate after 3 days at 30%. B) As A, but cells were also treated with 0.03% MMS during the exposure to 0.2M HU for 1-3 h. Colony scoring was carried out in triplicate after 3 days at 30°C. C) Serial dilution (10x) drop-assay on YPAD with indicated level of HU showing *rad18Δ* combined with *pol31*-W417 and *pol32Δ* mutants. The *rad18Δ* mutant on its own is sensitive to 20mM HU and the double mutant *pol32Δ rad18Δ* is synthetic lethal, yet *pol31-W417* is still able to suppress lethality in the triple mutant, *rad18Δ pol32Δ pol31-W417*. D) As C) but with *rev3Δ* mutant. The *rev3Δ* mutant on its own is not sensitive to 25 mM HU, yet pol31-W417 can -suppress the growth defects of *pol32Δ* on HU in the absence of *rev3Δ* (triple mutant *rev3Δ pol32Δ pol31-W417*). E) As C), but with *rad5Δ* mutant. The *rad5Δ* mutant on its own is not sensitive to HU, but the double mutant *pol32Δ rad5Δ* is synthetic lethal on 20mM HU. *pol31-W417* can still suppress the growth defect on HU in the triple mutant *rad5Δ pol32Δ pol31-W417*. F) As C), but with *mms2Δ* mutant. The *mms2Δ* mutant on its own is not sensitive to 25mM HU, yet shows additivity when combined with *pol32Δ* The *pol31-W417* allele can still suppress the growth defects on HU in the triple mutant *mms2Δ pol32Δ pol31-W417*.

We then returned to the drop assay, which allows us to compare single, double and triple mutants, in order to identify the pathways implicated in the suppression of *pol32Δ* deficiency, as done above on MMS. We find that *rad18Δ* cells, like *pol32Δ*, are sensitivity to growth on low level HU (Figure 4C). The suppression of *pol32Δ* by the *pol31-W417* allele, however, was largely independent of Rad18, and also independent of Rev3, Rad5, and Mms2 (Figure 4C-F). This is not surprising, yet it underscores the importance of a unique pathway of Polδ-dependent fork recovery, that is independent of PRRTLS and error-free lesion bypass.

### DNA polymerase Chromatin Immunoprecipitation (ChIP) at stalled replication forks

*We* have previously published methods for the quantitative monitoring of DNA Polα and Polε at replication forks stalled by 0.2M HU (Cobb, Bjergbaek et al., 2003, Cobb, Schleker et al., 2005). Since the collapse of the replication fork can be monitored by the loss or displacement of DNA polymerases near origins, we examined the impact of *pol31-T415* mutation and Pol32 ablation on DNA polymerase stability, and asked whether the suppression of *pol32Δ’s* HU sensitivity by *pol31*-W417 might act through the stabilization of DNA Polα and Polε at the replication fork.

We monitored the presence of Myc-tagged Pol1 and Pol2, the catalytic subunits of DNA Polα and Polε, at an early firing origin, and normalized polymerase abundance at 2, 4, 6, or 10 kb from the origin of replication, to its level at 14 kb, which is halfway to the next origin. *MATa* strains carrying epitope-tagged DNA polymerase Polα or DNA polymerase Polε were synchronized by α-factor and released synchronously into 0.2M HU, which arrests forks within 5 kb of initiation. In wild-type strains the polymerases are detected within 4-5 kb of the origin, for at least 1h on HU. In mutants such as *sgs1Δ, mrc1Δ or mec1Δ*, the polymerases are significantly less stable(Cobb et al., 2003, Cobb et al., 2005). Similar ChIP analyses show that in a *pol32Δ* mutant, i.e. in cells lacking Pol32, Polα (Figure 5A) as well as Polε (Figure 5B) are de-stabilized at a stalled replication fork. We concluded that the third subunit of DNA polymerase Polδ (Pol32) plays a role either directly or indirectly, in stabilizing polymerases other than Polδ at stalled replication forks.

**Figure 5:**
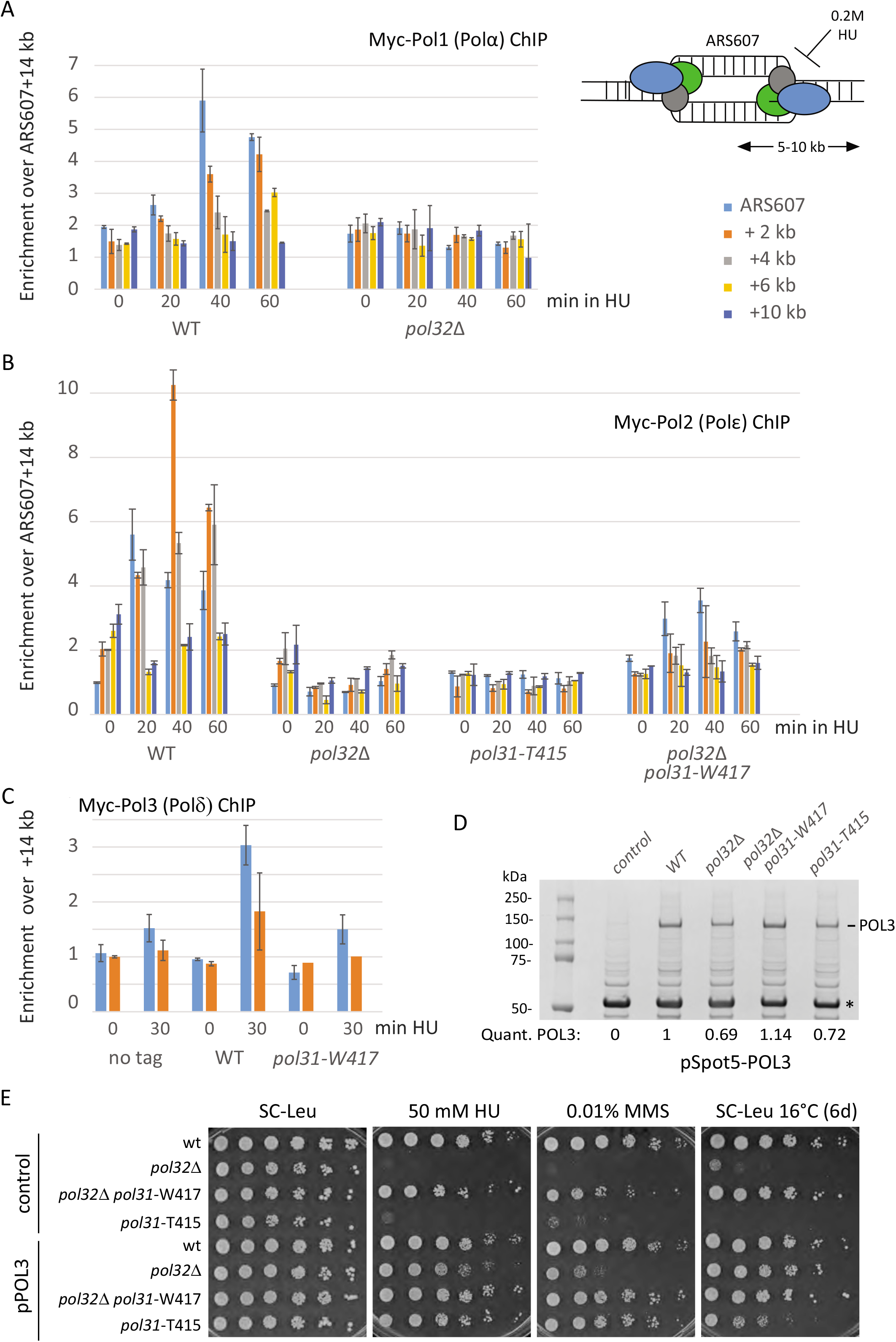
DNA polymerase binding at HU stalled replication forks is sensitive to*pol32Δ*. A) In wild-type and in *pol32Δ* cells, Pol1, the catalytic subunit of Pol, is destabilized at a stalled replication fork. Myc-tagged Pol1 is found enriched at stalled replication fork near early firing ARS607 on 0.2M HU, as demonstrated by Chromatin IP for (see Materials and Methods). Probes for qPCR are color-coded and are found at the indicated distances from ARS607, an efficient early firing origin on Chr VI. ChIP was performed 3 times and qPCR for the indicated probes was performed in triplicate for each biological replicate. Values are normalized to the signal at ARS607 + 14kb, which is halfway to the next origin of replication. B) As A, but for Myc-tagged Pol2, the catalytic subunit of Pole. Identical ChIP assays were also performed in strains bearing the *pol31-T415* and double *pol32Δ pol31-W417* mutations. The T415 mutation disrupts Pole enrichment, as does *pol32Δ* while *pol31-W417* partially compensates for the stabilization of Pole. C) As A, but for Myc-tagged Pol3, the catalytic subunit of Polδ. The polymerase is detected at stalled forks in wild-type and *pol31-W417* backgrounds. We were unable to generate Myc-tagged Pol3 in in *pol32Δ* and *pol31-T415* mutants, suggesting that the combination is synthetic lethal, even without HU. qPCR is shown only for ARS607 and +2kb, as these showed the strongest enrichment over nontagged Pol3 and the +14kb qPCR signal. Color-coded as A. D) Pull-down of SPOT-Pol3 shows that it is destabilized in the *pol32Δ* or *pol31-T415* background, while the double *pol32Δ pol31-W417* mutant restores SPOT-Pol3 levels. Cells with the indicated genetic background expressing N-terminally tagged SPOT-POL3 or control (SPOT peptide alone) from a plasmid, were grown to OD600 = 0.8 (exponential growth) in SC-LEU. Equal numbers of cells from each culture were lysed and the total cell lysate was incubated with an excess of SPOT-Trap Magnetic Agarose beads for 2h at 4°C. The SPOT-tagged Pol3 was recovered by IP, was denatured and an equal amount of each sample was loaded on 4-20 % SDS gel. The gel was stained with Instant Blue dye. SPOT-Pol3 is indicated and quantified by scanning, normalized to total protein in each lane. * indicates IgG. E) Serial dilution (lOx) drop assays on the indicated selective media was performed for strains with the indicated genotype bearing either a control plasmid (expressing SPOT tag alone) or the N-terminally tagged SPOT-Pol3 protein. Plates contained the indicated concentration of MMS or HU, or contained neither inhibitor and were simply incubated at 16°C for 6 days. All other plates were incubated for 3 days at 30°C prior to imaging. An extra copy of POL3 suppresses the MMS, HU and cold-sensitivity of *pol32Δ* and *pol31*-T415.

We next examined the effects of the *pol31* mutants on polymerase stability. ChIP analyses confirmed that Polε was destabilized at stalled replication forks by the pol31-T415 mutation (Figure 5B). Moreover, the suppressor *pol31-W417* partially restored Pol2-Myc recovery at stalled fork in *pol32Δ* cells. Although fork stability is very sensitive to the loss of Mec1, the checkpoint (monitored as Rad53 phosphorylation) was intact in all these mutants (Figure 1D), and the loss of Mec1 target sites on Pol31 (S20 and S51) to either alanine or glutamate residues, did not confer HU-sensitivity. We do note that coupling the phosphomimic Pol31 S20/S51 DD with *pol32Δ* was synthetic lethal, while the S20/S51 AA mutant helped overcome replication stress (Supplemental Figure 3).

We were able to perform ChIP for a C-terminally Myc-tagged DNA Polδ is extremely inefficient, and we were unable to create a Pol3 itself on HU, in wild-type cells and in the *pol31*-W417 mutant (Figure 5C). The enrichments were low, however, compared to that monitored with Pol1 or Pol2 (2-3 fold). We attempted to introduce the C-terminal Myc tag onto Pol3 or onto Pol31 in *pol32Δ* cells, or in the pol31-T415 mutant. However, despite multiple attempts, it was impossible to C-terminally tag Pol3 in *pol32Δ* or pol31-T415 mutant strains. This suggests that the combination of *pol32Δ* with tags that slightly compromise Polδ activity, is lethal. Thus, we could not monitor Polδ at stalled forks by ChIP in the strains that we propose might affect holo-Polδ stability, as we had monitored Pole.tagging strain in *pol32Δ* background (data not shown).

In order to see if Pol3 levels were indeed compromised by *pol32Δ* and/or pol31-T415, we tagged Pol3 with a small N-terminal Spot5 tag (peptide PDRVRAVSHWSS, Chromotek™), which we reasoned might be less disruptive to the complex than the large C-terminal Myc array. The Spot5 tag was inserted into the genomic copy of *POL3* and the resulting protein was recovered from a yeast cell lysate by binding to an excess of Spot-trap™ antibody-bound to magnetic beads (Figure 5D). The level of Spot5-Pol3 recovered from 1D). Earlier work had shown that the Pol31 N-terminus is a target of Mec1 on HU (Hustedt et al., 2015), but the mutation of the phosphoacceptor sites (S20 and S51) to either alanines or glutamate residues, that ablate or mimic phosphorylation respectively, did not alone confer HU sensitivity. We do note that coupling the phosphomimic Pol31 S20/S51 DD with pol32*Δ* extracts was 70% of the level recovered in wt cells, and the combination of *pol32Δ* with *pol31-W417* suppressed the drop, nearly doubling it over the level found in *pol32Δ* cells. Like *pol32Δ*, the pol31-T415 mutant reduced the amount of Pol3 recovered (Figure 5D). To see if this change in Pol3 level was responsible for the HU-, MMS-or cold sensitivity of the *pol32Δ* mutant, we introduced a plasmid that expressed an additional copy of *POL3* from a plasmid (pSpot5-POL3). As expected, with a control plasmid *pol32Δ* and pol31-T415 mutant strains showed HU-, MMS- and cold-sensitive growth, while the presence of *pol31-W417* suppressed the sensitivity of *pol32Δ* for all phenotypes (Figure 5E). Interestingly, raising the level of Pol3 in the pol31-T415 mutant also suppressed its slow growth under fork stalling conditions (Figure 5E). We conclude that Pol31 and Pol32 stabilize holo-Polδ in the context of a stalled or collapsed forks. Given that Pol3 levels drop in *pol32Δ* and that this can be suppressed by the *pol31-W417* mutation which increases affinity between Pol31 and the Pol3-CTD, we conclude that Pol3 levels are critical for the survival of fork stress conditions. Suppression of *pol32Δ’s* sensitivity to alkylating damage may also involve Pol31-mediated stabilization of the TLS polymerase, Polζ (Rev3), although we did not see any impact of Rev3 on HU-induced replication stress.

One pathway for fork recovery after HU induced fork collapse is Break-induced Replication (BIR), a recombination-dependent pathway that is largely dependent on Pol32 (Lydeard et al., 2007). Using a well characterized BIR assay in which an induced HO-endonuclease double-strand break triggers exchange between two unrelated chromosomes, we could reproduce the fact that successful BIR requires Pol32 (Lydeard et al., 2007). Interestingly, in this assay the pol31-T415 and *pol31-W417* mutants were both less efficient than wild-type cells, yet the pol31-T415 allele, which in other assays shows sensitivities equal to *pol32Δ*, was far less compromised for BIR than cells lacking Pol32 (Supplemental figure 4). Consistent with the interpretation that the role played by Polδ in BIR is distinct from that monitored on MMS or acute exposure to HU, we find that the *pol31-W417* mutant is unable to significantly suppress the loss of BIR by ablation of Pol32 (Supplemental Figure 4). The mechanism of initiation of BIR through a DSB and a collapsed fork may differentiate the role of Pol32 and Pol31 in BIR or template switch synthesis. We conclude that Pol32 plays an important role in the resumption of replication and/or maintenance of polymerase stability when replication forks stall or collapse, and Pol31 can compensate for some, but not all of the *pol32Δ* associated defects by maintaining functional levels of Pol3.

## Discussion

### DNA polymerase δ in DNA replication and repair

Many of the pathways that control replication fork recovery under control of Mec1/ATR make use of the lagging strand polymerase, DNA Pol δ (Hubscher & Maga, 2011, Maga & Hubscher, 2008). In budding yeast Polδ is a heterotrimeric enzyme consisting of the catalytic subunit Pol3 (human POLD1/p125), Pol31 (POLD2/p50), and Pol32 (POLD3/p66). The human enzyme also contains a smaller regulatory subunit POLD4/p12. The first and the second subunits of yeast Polδ are encoded by the essential genes *POL3/CDC2* and *POL31/HYS2/SDP5*, respectively, but the third subunit is encoded by the non-essential gene, *POL32*. The DNA polymerase processivity factor, PCNA interacts through the multiple PIP sites in Pol δ, with strong interaction through the C-terminus of Pol32 and/or Pol3 itself (Fuchs et al., in press),(Acharya et al., 2011, Johansson et al., 2004).

An increasing body of evidence implicates the Pol31 and Pol32 subunits of Polδ in DNA repair, particular at the replication fork or in PRR pathways for the repair of alkylated bases. Consistently, Pol31 and Pol32 were found as core subunits of the DNA Polζ, which mediates error-prone Ţranslesion Synthesis (TLS) (Johnson et al., 2012, Makarova et al., 2012). Intriguingly, a recent study in chicken DT40 cells found that POLD3 has an additive contribution to TLS independent of Polζ (Hirota et al., 2015). It was later shown that inactivating the proofreading activity of POLD1 restores TLS activity in POLD3-deficient chicken DT40 cells, strongly implicating Polδ is able to contribute to TLS (Hirota et al., 2016). A similar finding in budding yeast demonstrated that Pol32 contributes to TLS independent of Polζ (Siebler et al., 2014). Consistent with an important function of Pol32 in dealing with DNA fork associated damage, we confirm that cells lacking Pol32 show cold-, HU- and MMS-sensitivity. Importantly, in all assays on MMS, the *pol31-T415* mutant behaves very much like the *pol32* null allele, impairing the resumption of replication after replication fork stalling on MMS, while the *pol31-W417* mutant efficiently suppresses the MMS sensitivity of *pol32Δ We* attribute this to the altered interaction of Pol31 with Pol3 which in turn stabilizes Polδ activity.

The suppression of *pol32Δ* sensitivity to HU and the defects of the pol31-T415 mutant, are similar to that observed on MMS, even though the pathways of recovery are very different. The impact of the mutations was monitored not only in drop-assays, but as recovery from acute exposure to 0.2M HU, which blocks replication forks genome-wide. Finally ChIP analyses showed that the absence of Pol32 destabilizes DNA Polε and Polα at stalled replication forks, as does the cold-sensitive mutant *pol31*-T415. This implicates the smaller subunits of Polδ in general replisome stability, which may be an indirect effect of stabilization of Pol3. Finally, the Pol32 subunit is known to be required for the efficient completion of Break-Induced Replication (BIR), a recovery pathway that allows restart of stalled or broken replication forks through invading a template sequence with homology to one end of the DSB. We find that the *pol31* mutations studied here are only partially deficient in BIR and the W417 suppressor allele does not efficiently suppress the loss of BIR provoked by ablation of Pol32. In other studies, Pol32 was shown to be required for both Rad51-dependent and independent survivors in the absence of telomerase (Lydeard et al., 2007).

In human and *Drosophila* cells it has been shown that the loss of POLD3 or POLD2 leads to the degradation of the other subunits of holo-Pol δ (reviewed in Fuchs et al., in press). We show a similar drop in Pol3 levels upon loss of Pol32, and/or introduction of the mutation pol31-T415. The instability of Pol3 was restored by combining *pol32Δ* with *pol31-W417*. Exactly the same sensitivity and suppression is observed by the expression of an ectopic copy of wild-type *POL3* (Figure 5E). This argues that the stability of the holo-Polδ complex plays a crucial role both in dealing with alkylation by PRR, and in the resumption of replication and/or maintenance of the replisome when replication forks are stalled on HU.

These observations are relevant with respect to the overexpression and upregulation of POLD2 and POLD3 in human cancers, as it suggests that their upregulation essentially upregulates and stabilizes POLD1. These observations also suggest that an inhibition of Polδ in cancer cells might sensitize tumor cells to survival of damage. Under the persistent replication stress that is found in transformed cells, we propose that the role of POLD2/Pol31 is to stabilize POLD1/Pol3. This is supported by our data showing that the loss of Pol32 can be compensated by the *pol31-W417* mutant which increases affinity between Pol31 and Pol3. Under PRR conditions the same mutation may similarly confer a more stable interaction with the translesion synthesis polymerase, Rev3. The switch between polymerases may be controlled by the checkpoint kinase Mec1-Ddc2, given that Pol31 is a target of Mec1 phosphorylation specifically in S phase. The S/T to D mutation of Mec1 target sites in Pol31 is synthetic lethal with *pol32Δ*, while the S/T to A mutations slightly suppress *pol32Δ* sensitivities to replication stress.

Based on the crystal structure of the human orthologue of Pol31-Pol32_N-term_ (Baranovskiy et al., 2008), we built a homology model of Pol31, to which we mapped the positions of these novel *pol31* mutations. The cs pol31-T415 and the *pol31-W417* suppressor map to a highly conserved, solvent exposed loop on the surface of Pol31 far from the Pol32 and PCNA binding interfaces (Fig. 1C). We examined the involvement of this exposed loop in protein-protein interactions, by performing yeast two-hybrid and pull-down assays with the Pol3 C-terminal domain (CTD). Consistent with our damage-survival data, pol31-T415 results in the loss of interaction between Pol31 and Pol3-CTD while *pol31-W417* enhances it (Figure 2). These results demonstrate clearly the impact of Pol3-Pol31 interaction on the stability and function of Pol δ. Intriguingly, the pol31-T415 and *pol31-W417* mutations also respectively disrupt and enhance the interaction of Pol31 with the CTD of the TLS polymerase, Rev3 (Pol ζ).

The solvent exposed loop of Pol31 containing residues T415 and W 417 is necessary to bind the Fe-S cluster CysB of Pol3, as well as the C-terminal Zn finger of Rev3. Thus, the molecular mechanism of Polδ destabilization and/or stabilization under conditions of stress, can be traced largely to this interaction domain. Nonetheless, the pol31-W417 mutant does not compensate for the impact of *pol32Δ* on BIR, as the *pol31-W417* suppressor allele does not restore efficient BIR in *pol32Δ*. It remains to be seen what additional roles Pol32 plays in this event beyond the stable recruitment of Pol3. It has been suggested that the role of Pol32 in BIR is primarily through the binding of its N-terminus to Pol31 (Lydeard et al., 2007).

Surprisingly, and in contrast to previous studies which reported suppression of the defects of the Polδ mutants by deletion of *RAD18*, such deletions aggravated the temperature sensitivity conferred our *pol31* alleles. Moreover, the pol31-W417 allele was not able to suppress *pol32Δ* in the absence of Rad18 for survival on HU. This suggests that Rad6-Rad18 may play a role in the efficient restoration or resumption of replication at stalled forks. Probing this and its relevance for Polδ activity, will be the focus of future studies.

## Acknowledgements

We thank the Haber lab for useful strains and Nicole Hustedt for the generation of the phosphosite mutants of Pol31. We are grateful to the Gasser lab for helpful suggestions to this work. We also acknowledge the generous financial support of the Swiss National Science Foundation grant number 31003A-176286 to S.M.G and the Swiss Cancer League grant 4167-02-2017 to SMG and ND. We thank the Novartis Research Foundation for continued core support of the FMI.

## Materials and Methods

### Yeast culture, strains, plasmids

The yeast strains and plasmids are described in Tables S1 and S2. Mutant *pol31* yeast strains were prepared by plasmid shuffling. If not stated otherwise cells were cultured at 30°C in YPAD medium using standard procedures, unless otherwise indicated.

### Yeast Two Hybrid

For yeast two hybrid analysis, WT and mutants of Pol31, and the full-length Rev3 were fused to the lexA DNA binding domain of pGAL-LexA (Bait). The WT and mutants of Pol31, Pol32, PCNA and the C-terminal CysA and CysB domains of Pol3 were fused to the B42 transcription activation domain of pJG4-7 (Prey). Liquid β-galactosidase assays were performed as previously described (Hegnauer et al., 2012) using the WT yeast strain (GA-1981) or *pol32Δ* mutant (GA-6292) containing the lacZ reporter pSH18-34, the bait and the prey. Exponentially growing, glucose-depleted cells were exposed to 2% galactose for 6 h to induce the fusion proteins, and protein-protein interactions were detected by the quantitative β-galactosidase assay of cell extracts. Three to four independent transformants were analysed for each interaction. Expression of all the fusion proteins was confirmed by western blot analysis (data not shown).

### Pull-down assays

Lysis buffer was supplemented with protease inhibitors (Complete protease inhibitors, Roche), Yeast cell extract containing over-expressed HA-Pol31 derivatives (B42 / WT / T415 / W417, plasmids #1493, #3062, #3063, #3064) were added to *E*. co//-expressed MBP-Pol3-CTD bound to Amylose beads: CysA + CysB (Pol3 _991-1097_) or only CysB (Pol3 _1029-1097_), See Figure S1 for a general outline of the pull-down assay. The amount of used Pol3 CysA + CysB domains were detected by UV-imaged Criterion gels (BioRad) and the binding of Pol31 to Pol3-CTD was analysed by Western blot using anti-HA antibody.

Proteins were coupled to beads in the lysis buffer (50 mM HEPES pH7.5, 20 mM NaCl, 1 mM EDTA, 0.1% Triton X-100, protease inhibitor cocktail (Roche)) for 1.5h at room temperature, then washed three times with lysis buffer before use. Exponentially growing yeast cells of the indicated genotype (~l × 10^7^ cells/ml, 200 ml) were pelleted and washed once with ice-cold PBS. Cell pellet was resuspended in 0.8 ml lysis buffer and the cell lysate was prepared by beat beating with Zerconia beads, 6.5 Hz, 60 sec, 4 times at 4°C. Cell lysate was clarified by centrifugation 12000 x g, 5 min at 4°C. 150 μl of total cell lysate was incubated with MBP-Pol31 bound to Agarose beads for 1.5h at 4°C with constant rotation, and beads were washed three times with excess lysis buffer containing 0.1% TritonX-100 and 20 mM NaCl at 4°C. Pull-down proteins were eluted from the beads and the protein sample was denatured and boiled in 1× NuPAGE sample buffer, and analysed by NuPAGE followed by silver staining and Western blot.

### Drop assay, cell recovery assay, checkpoint activation

All methods for checkpoint activation by phospho-shift analysis of Rad53 phosphorylation, test for cell sensitivity to DNA damaging agents by drop assay, cell recovery assay from MMS after α-factor synchronization were performed as described in (Hustedt et al., 2015).

To monitor Rad53 activation by Western blot, exponentially growing cells (GA-1981, GA-5321, GA-5076, GA-5324, GA-1761, GA-2056) were synchronized in G1 with α-factor and released into 0.2 M HU, or 0.03% MMS containing YPAD media at 30°C for 0.5 - 1 h. Cold-sensitivity of pol31 mutants were released into YPAD media at 16°C and grown for 1 h at 16°C. For Western Blot analysis, lysates were prepared using silica beads and urea buffer and subjected to 7.5 % SDS-PAGE, transferred onto PVDF membrane and detected with anti-Rad53 antibody (Santa Cruz sc-6749).

### ChIP analysis

ChIP experiments were performed as described (Cobb et al., 2003, Cobb et al., 2005). Monoclonal anti-HA (F-7, Santa Cruz) was used to precipitate HA-tagged DNA pol α, and anti-Myc (9E10) to precipitate Myc-tagged Ddc2. BSA-saturated Dynabeads were coupled to the monoclonal antibodies and incubated with the cell extract for 2 h at 4 °C. BSA-coupled Dynabeads without each strain are averaged over three independent experiments with real-time PCR performed in duplicate (error bars indicate standard error of the mean). Absolute fold enrichment at ARS607 was calculated as follows. For each time point the signal from the antibody-coupled Dynabeads was divided by the signal from the BSA-coated Dynabeads, after both signals were first normalized to the signal from the input DNA. Finally, the relative enrichment for ARS607 was obtained by normalizing the absolute enrichment at the indicated probe near ARS607 for each timepoint to the absolute enrichment at a locus 14 kb away from ARS607.

### Survival and drop assays

For liquid survival assays, overnight cultures were diluted to OD_600_ = 0.15 and grown for 3 h, then synchronized with α-factor in G1 and released into 0.2 M HU containing YPAD. After the indicated time points relevant dilutions were plated onto fresh YPAD plates and colonies were counted after 3 to 4 days. Survival is defined as the fraction of indicated doses compared to the untreated control (0h) normalized to the survival of WT cells for each time point. For drop tests, overnight cultures were diluted to a starting density of OD_600_ = 0.5 and 2 μl drops with 4 x 10-fold dilutions were plated on YPD or the appropriate selective medium containing freshly diluted concentrations of MMS or HU as indicated.

### BIR assay

Mainly following the published protocols in Anand et al (2014) and Lydeard et al (2014), we used the2010 (Anand, Tsaponina et al., 2014, Lydeard, Lipkin-Moore et al., 2010). BIR-tester strain, GA-8997, in which the *CAN1* gene is replaced by *URA3* and the *HPH* by clonNAT (gift of J. Haber’s lab). A galactose inducible HO endonuclease generates a DSB within the *UR3-NAT* locus on Chr V. Repair by BIR of the DSB results in URA+ and clonNAT-sensitive viable cells. In GA8997 the donor *5’-UR3* is 33kb away from Telo-V-R and the Recipient *3’-RA3* is 39kb away from Telo-IX-R. All strains from –80°C were directly plated on YPAD + clonNAT plates to use only cells with intact HO-sites. Then sc-cultures were grown in YPA-2% Raffinose for 6hrs to 10^7^ cells/ml and 10-fold serial dilutions were plated each on YPAD (to get the total cell count), on SC-ura (to count for URA+-background) and on YPA-Galactose to induce the HO endonuclease. Cell viability after HO-endonuclease-induction was derived by dividing the number of colony forming units (CFU) on YEP-Galactose by that on YEPD. Rate of BIR-repair was determined by URA+ CFU divided by YEPD CFU multiplied to the number of URA+ CFU divided by YEP-Gal CFU.

A PCR-based 9-Myc epitope tagging method was used to tag *POL1, POL2, POL31* and *POL3* genes by chromosomal integration of PCR-amplified cassette in the KSC106 yeast background used for genetic analyses. The correct integration of the cassette has been verified by PCR as well as checking the expression of the Myc-tag by Western blotting. To see if the hypersensitivity of the isolated ts/cs *pol31* mutants reflects polymerase destabilization, we epitope tagged *POL3* with the Nterminal Spot tag (). These cells were viable and the tag did not compromise Polδ function, whereas 9-Myc tagged Pol3 was lethal when combined with *pol32Δ*

### Homology modeling of the Pol31-Pol32N complex

HHPRED (Soding et al., 2005) was used to search the Protein Data Bank (www.rcsb.org) for Pol31 and Pol32 orthologues of known structure. For both Pol31 and Pol32, the crystal structure of the human P50-P66(N) complex (PDB 3E0J, (Baranovskiy et al., 2008)) was the top hit which gave high scores for Pol31 matching P50 (probability=100%, E-value=1.2e-66, identities=32%, Pol31 residues 19-486) and Pol32 matching P66(N) (probability=99.93, E-value=3.4e-26, identity=13%, Pol32N residues 1-127). HHPRED pairwise sequence alignments for Pol31/P50 and Pol32/P66(N) were locally corrected, then merged and used for modeling the heterodimeric complex using the modeller software (Sali & Blundell, 1993) together with the 3E0J crystal structure as 3D template (chains A, B). Pol31 loops longer than 10 residues without structural information in the 3D template were excluded from modeling calculations. 250 models for the Pol31-Pol32N complex were generated and the best model was chosen according to the lowest values for the modeller objective function and the quality of the Ramachandran plot.

## Supplemental figures

**Figure Supplemental 1.**
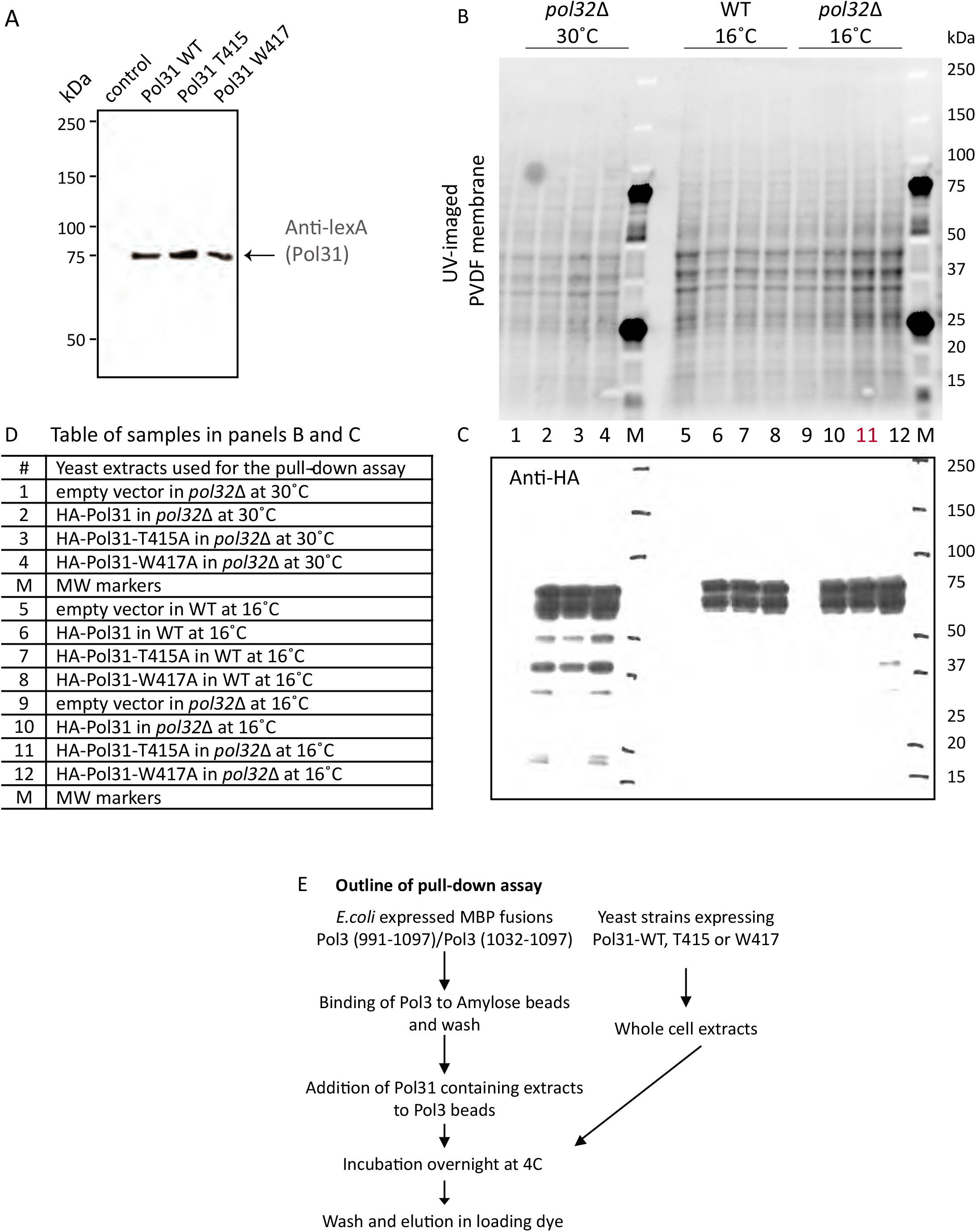
Protein expression in 2 hybrid assays and pull-down fusion constructs. A) Western blot showing equal expression of Pol31 wild-type and the indicated mutants used for the yeast two-hybrid assays shown in Figure 2. Blot is probed with anti-LexA. B) Amido black staining for protein extracts used for the pull-down assay in Figure 2E. C) Western blot of PVDF membrane showing equal level expression of HA-Pol31 under all conditions of pull-down in all mutants used (see panel D). D) Table of samples used for the pull-down assay. E) Scheme of the pull-down assay shown in Figure 2.

**Figure Supplemental 2.**
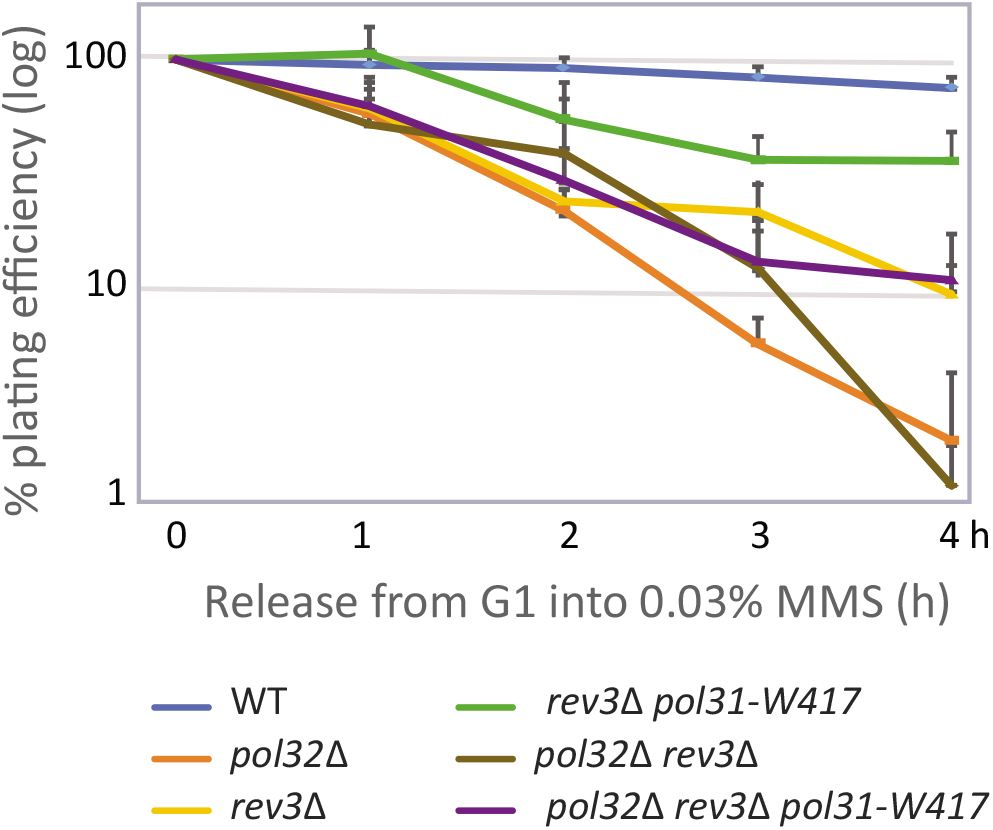
Colony outgrowth assay after exposure to MMS. The *pol31-W417* suppression of *pol32Δ* sensitivity to MMS is independent of the TLS polymerase Rev3. A survival assay was performed as in Figure 3A. Quantitative colony counting showing that the pol31-W417 mutation suppresses the growth defects of *pol32Δ* in liquid culture on 0.03% MMS, even in the absence of Rev3. Cells were G1-arrested by α-factor, released into 0.03% MMS, sampled at indicated time-points and plated on YPAD plates in triplicate. Colonies plated in triplicate are scored after 3 days.

**Figure Supplemental 3.**
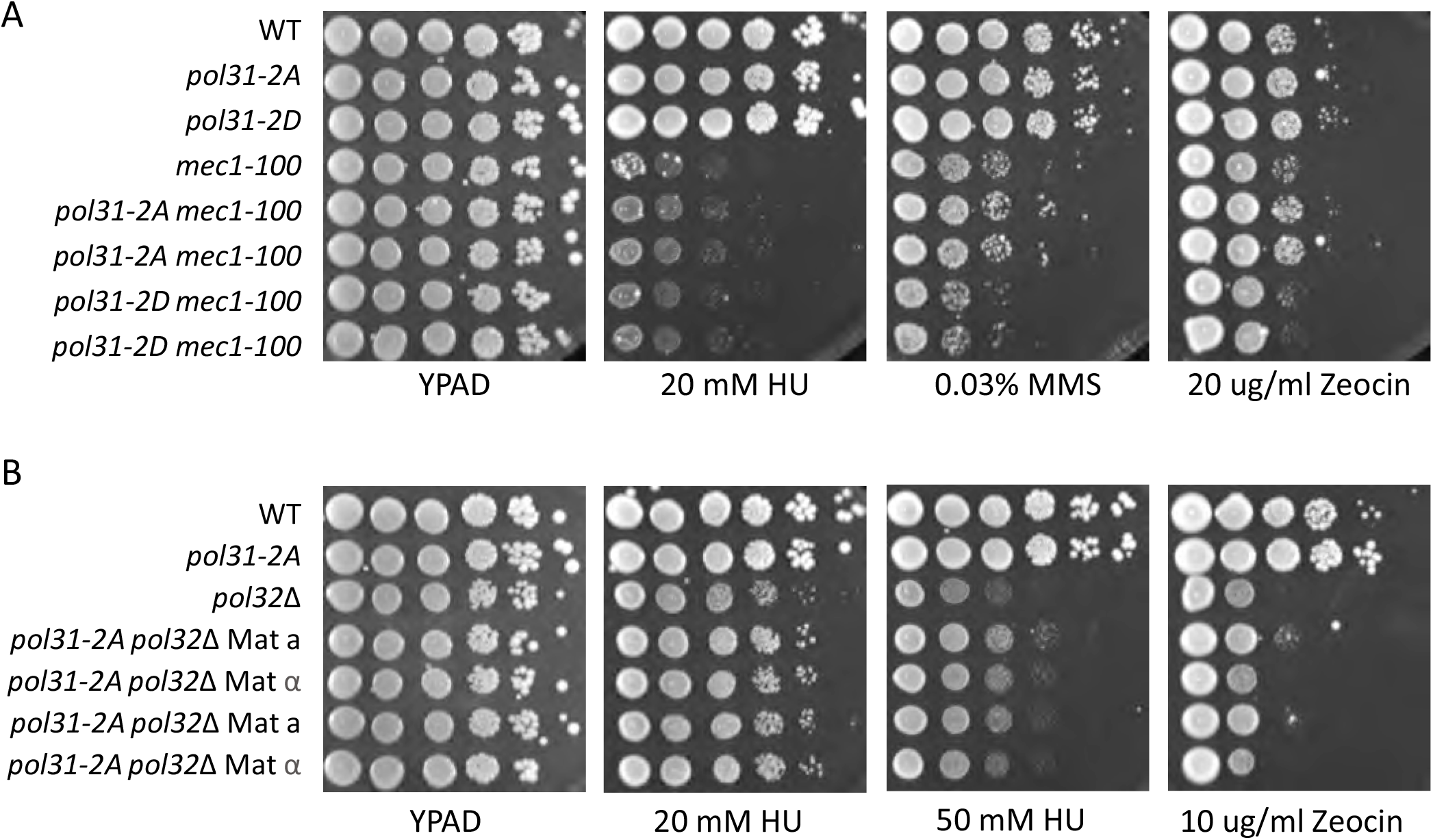
Mec1 phosphoacceptor sites on Pol31 do not affect survival of DNA damage. Pol31 is a target of Mec1 on HU on the residues S20Q and S51Q (Hustedt et al., 2015). We mutated both phospho-acceptor residues either to alanine *(pol31-2A)* or glutamic acid *(pol31-2D)*. Serial dilution (lOx) series on appropriate media containing the indicated reagents shows that the mutants do not show hypersensitivity to HU, MMS nor to Zeocin on their own. The mutations neither suppress nor show additivity with *mec1-100*, which compromises the phosphorylation (Hustedt et al., 2015). The combination of *pol31-2A pol32Δ* slightly suppresses MMS sensitivity of *pol32Δ* by itself, yet the combination of *pol31-2D pol32Δ* was shown to be synthetic lethal (data not shown). Since the pol31-2D phenotype is manifest in the absence of Pol32, the phosphorylation of Pol31 S20 and S51 may affect the binding of another factor that is necessary to stabilize Pol3 activity.

**Figure Supplemental 4.**
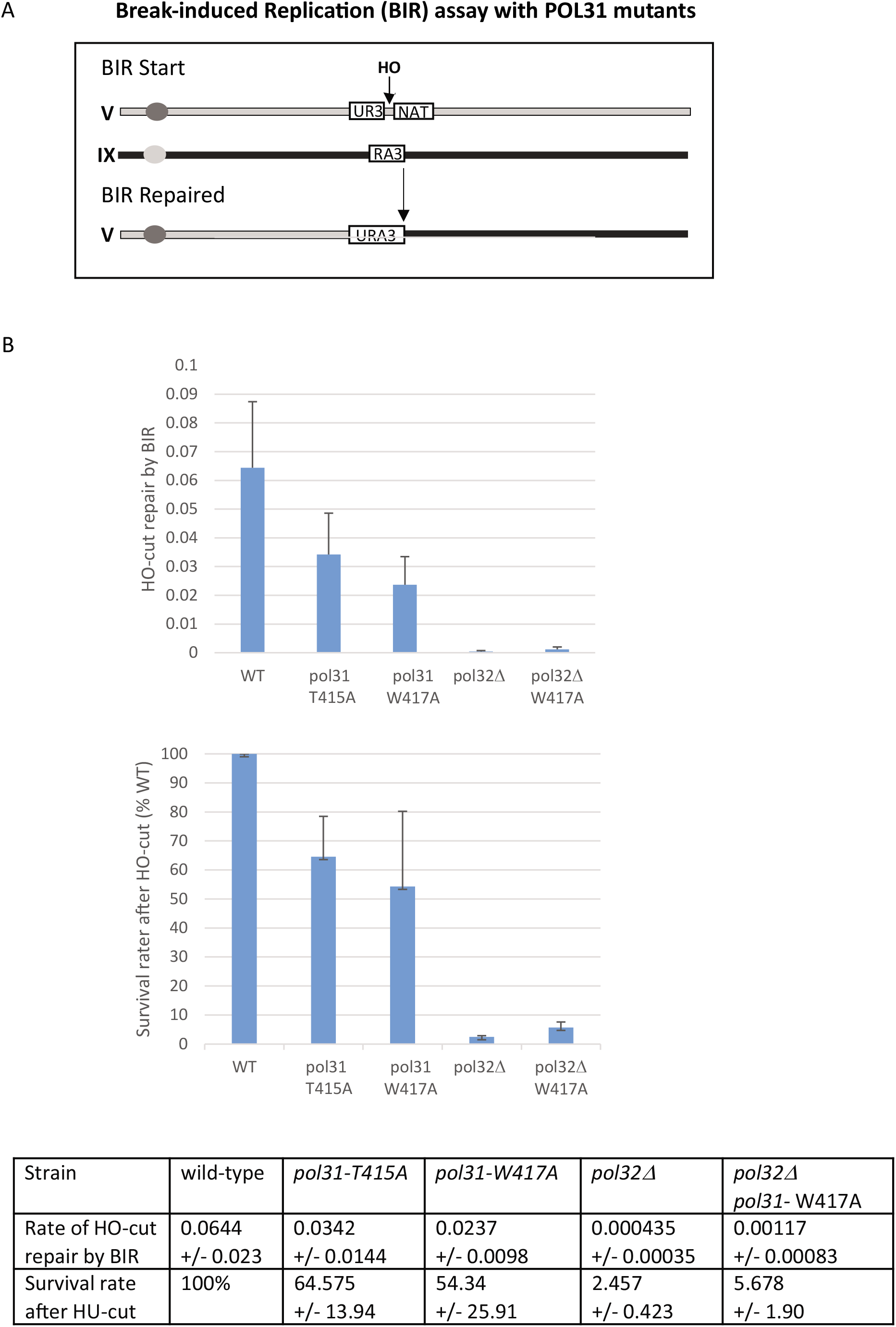
Break-induced replication assay: *pol31-W417* cannot replace *pol32Δ* for BIR. A) The *CAN1* gene is replaced by *URA3* and the *HPH* by clonNAT in GA-8997 (gift of J. Haber). After HO-break induction, a homology-driven repair event between the two chromosomes is mediated by BIR, restoring uracil prototrophy and clonNAT-sensitive viable colonies. In the GA-8997 strain the donor 5’-UR3 is 33kb away from Telo-V-L and the recipient 3’-RA3 is 39kb away from Telo-IX-L. B) BIR efficiency was scored in GA-8997 which is wild-type for all subunits of Polδ, and in the same background bearing the indicated *pol31* and *pol32Δ* mutations. Each assay was quantified in triplicate and each assay was replicated at least 3 times. BIR is strongly affected by *pol32Δ*, and is reduced by both pol31-T415A and pol31-W417A mutations. Importantly, only minor suppression of *pol32Δ* by *pol31-W417A* can be detected by an HO-induced BIR assay. We confirmed that HO cut efficiency was comparable in all strains, and repair efficiency of induced HO cleavage was normalized cut efficiency.

**Table S1:** Yeast strains and plasmids used in this study

| Strain | Genotype | Plasmid | Source |
| --- | --- | --- | --- |
| GA-4619 | (KSC106) *MATa, ade1, his2, trp1, ura3, leu2* |  | (K. Sugimoto) |
| GA-4620 | GA-4619 with *pol31::kanMX4* | pRS316-POL31 | (Davoodi et al., 2006) |
| GA-9784 | GA-4619 with *pol31::kanMX4, pol31-EF463AA* | pPOL31-EF463 | this study |
| GA-9782 | GA-4619 with *pol31::KanMX4, pol31-W417A* | pPOL31-W417 | this study |
| GA-4761 | GA-4619 with *pol32::hphMX* |  | this study |
| GA-4639 | GA-4619 with *pol31::kanMX4, pol32::hphMX, pol31-EF463AA* | pPOL31-EF463 | this study |
| GA-4636 | GA-4619 with *pol31::kanMX4, pol32::hphMX, pol31-W417A* | pPOL31-W417 | this study |
| GA-4630 | GA-4619 with *pol31::kanMX4, pol31-T415A* | pPOL31-T415 | this study |
| GA-1981 | *MATa, ade2-1, trp1-1, his3-11, -15, ura3-1, leu2-3, -112, can1-100 (W303), RAD5+* |  | (H.L. Klein) |
| GA-6292 | GA-1981 with *Pol2-13Myc-G418, pol32::natMX* |  | this study |
| GA-4780 | GA-4619 with *pol31::KanMX4, rev3::LEU2* | pRS316-POL31 | this study |
| GA-9207 | GA-4619 with *pol31::kanMX4, pol31-W417A, rev3::natMX, pol32::hphMX,* | pPOL31-W417 | this study |
| GA-9644 | GA-4619 with *pol32::hphMX4, rev3::natMX* |  | this study |
| GA-9780 | GA-4619 with *pol31::KanMX4, rev3::LEU2, pol31-W417A* | pPOL31-W417 | this study |
| GA-9664 | GA-4619 with *pol31::KanMX4, rad18::natMX* | pRS316-POL31 | this study |
| GA-9665 | GA-4619 with *pol32::hphMX, rad18::natMX* |  | this study |
| GA-9666 | GA-4619 with *pol31::KanMX4, pol31-W417A, pol32::hphMX, rad18::natMX* | pPOL31-W417 | this study |
| GA-4817 | GA-4619 with *rad5::hphMX* |  | this study |
| GA-9779 | GA-4619 with *pol31::kanMX, pol32::natMX, rad5::hphMX* | pRS316-POL31 | this study |
| GA-9778 | GA-4619 with *pol31::kanMX, pol31-W417A, rad5::hphMX* | pPOL31-W417 | this study |
| GA-9781 | GA-4619 with *pol31::kanMX, pol31-W417A, pol32::natMX, rad5::hphMX* | pPOL31-W417 | this study |
| GA-7566 | GA-1981 with *hphMX-pol31-2D* |  | this study |
| GA-7600 | GA-1981 with *hphMX-pol31-2A* |  | this study |
| GA-8997 | *MATa::DEL. HOcs::hisG ura3D851 trp1DEL.63 leu2DEL::KAN hmlDEL::hisG hmrDEL::ADE3 ade3::GAL::HO can1DEL::UR:: HOcs::NAT, RA3::TRP1 (at SUC2 locus in Chr IX)* |  | (J. Haber) |
| GA-9133 | GA-8997 with *pol32::hphMX* |  | this study |
| GA-9137 | GA-8997 with *pol31-T415A* |  | this study |
| GA-9138 | GA-8997 with *pol31-W417A* |  | this study |
| GA-4821 | GA-4619 with *mms2::hphMX4* |  | this study |
| GA-9893 | GA-4619 with *mms2::hphMX4 pol32::hphMX4* |  | this study |
| GA-9891 | GA-4619 *MATα* with *mms2::hphMX4 pol32::hphMX4 pol31::kanMX4 pol31-W417A* | pPOL31-W417 | this study |
| GA-5050 | GA-1981 with *POL2-13Myc_KanMX* |  | this study |
| GA-6292 | GA-5050 with *Pol2-13Myc_KanMX, pol32::NAT* |  | this study |
| GA-10007 | GA-5050 with *Pol2-13Myc-G418, pol31::URA3* | pPOL31-W417 | this study |
| GA-10008 | GA-5050 with *Pol2-13Myc-G418, pol31::URA3* | pPOL31-T415 | this study |

**Table S2:** Plasmids used in this study

| Plasmid No. | Plasmid Name | Source |
| --- | --- | --- |
| # 2731 | *pBL331 (pRS314-POL31 (-139 to +2044, ARS CEN TRP1))* | (Gerik et al., 1998) |
|  | *pRS316-POL31 (-139 to +2044, ARS CEN URA3)* | (Davoodi et al., 2006) |
| # 3881 | *pPOL31-EF463 (-139 to +2044, ARS CEN TRP1)* | this study |
| # 2732 | *pPOL31-W417 (-139 to +2044, ARS CEN TRP1)* | this study |
| # 2736 | *pPOL31-T415 (-139 to +2044, ARS CEN TRP1)* | this study |
| # 359 | *pSH18-34 (2μ, URA3)* | (Golemis et al., 2011) |
| # 965 | *pGAL-LexA (2μ, HIS3)* | (Bjergbaek et al., 2005) |
| # 2686 | *pGAL-LexA-POL31 (aa1-487, 2μ, HIS3)* | this study |
| # 2856 | *pGAL-LexA-POL31-T415 (aa1-487, 2μ, HIS3)* | this study |
| # 2855 | *pGAL-LexA-POL31-W417 (aa1-487, 2μ, HIS3)* | this study |
| # 3816 | *pGAL-LexA-REV3 (aa1-1504, 2μ, HIS3)* | this study |
| # 1493 | *pJG47-HA (2μ, TRP1)* | this study |
| # 2693 | *pJG45-HA-POL3 (aa991-1097, 2μ, TRP1)* | this study |
| # 2694 | *pJG45-HA-POL3 (aa1032-1097, 2μ, TRP1)* | this study |
| # 3062 | *pJG47-HA-POL31 (aa1-487, 2μ, TRP1)* | this study |
| # 3063 | *pJG47-HA-POL31-T415 (aa1-487, 2μ, TRP1)* | this study |
| # 3064 | *pJG47-HA-POL31-W417 (aa1-487, 2μ, TRP1)* | this study |
| # 2949 | *pJG47-HA-PCNA (aa1-259, 2μ, TRP1)* | this study |
| # 3010 | *pJG47-HA-POL32 (aa1-350, 2μ, TRP1)* | this study |
| # 2905 | *pOPINM His-MBP-Pol3 (aa991-1097, T7-lacO, pUC-ori, Amp)* | this study |
| # 2906 | *pOPINM His-MBP-Pol3 (aa1032-1097, T7-lacO, pUC-ori, Amp)* | this study |
| # 3115 | *pRS406-pol31AQ (S20A, S51A, URA3)* | this study |
| # 3116 | *pRS406-pol31DQ (S20D, S51D, URA3)* | this study |
| # 3716 | *pRS305-POL31-T415 (aa1-487, LEU2)* | this study |
| # 3717 | *pRS305-POL31-W417 (aa1-487, LEU2)* | this study |
| #4072 | *pSPOT5 (CEN, LEU2)* | Chromotek |
| #4073 | *pSPOT5-POL3* | this study |

